# The origins of a novel butterfly wing patterning gene from within a family of conserved cell cycle regulators

**DOI:** 10.1101/016006

**Authors:** Nicola J. Nadeau, Carolina Pardo-Diaz, Annabel Whibley, Megan Supple, Richard Wallbank, Grace C. Wu, Luana Maroja, Laura Ferguson, Heather Hines, Camilo Salazar, Richard ffrench-Constant, Mathieu Joron, W. Owen McMillan, Chris D. Jiggins

## Abstract

A major challenge in evolutionary biology is to understand the origins of novel structures. The wing patterns of butterflies and moths are derived phenotypes unique to the Lepidoptera. Here we identify a gene that we name *poikilomousa* (*poik*), which regulates colour pattern switches in the mimetic *Heliconius* butterflies. Strong associations between phenotypic variation and DNA sequence variation are seen in three different *Heliconius* species, in addition to associations between gene expression and colour pattern. Colour pattern variants are also associated with differences in splicing of *poik* transcripts. *poik* is a member of the conserved fizzy family of cell cycle regulators. It belongs to a faster evolving subfamily, the closest functionally characterised orthologue being the *cortex* gene in *Drosophila,* a female germ-line specific protein involved in meiosis. *poik* appears to have adopted a novel function in the Lepidoptera and become a major target for natural selection acting on colour and pattern variation in this group.

## Introduction

The wings of butterflies and moths (Lepidoptera) are some of the most diverse and striking examples of evolutionary diversification. They are variously implicated in crypsis, warning colour and sexual selection. The patterns consist of arrays of coloured scales, which are a unique feature of the eponymous Lepidoptera. The evolution of these scales from sensory bristles is an example of developmental novelty that is amenable to study in the field and laboratory (1). Scale colours appear to be intimately linked to scale development, as scales of different colours develop at different rates and have different morphologies (1,2). The evolutionary innovation of coloured wing scales produced new adaptive potential. For example, eyespots on the wing increase survival by acting as startle patterns to drive off predators (3), while industrial melanism and its reversal in the peppered moth is one of the most striking examples of recent evolution (4). Divergence in wing patterning has also been implicated as a primary cause of speciation in *Heliconius* butterflies (5).

After a long history of ecological study, we are now starting to understand the genetic regulation of wing patterning. For example, the *doublesex* gene has recently been shown to control female limited mimetic polymorphism in the swallowtail *Papilio polytes* (6). However, other swallowtails have evolved sex-limited mimicry using different genetic loci (7,8). In contrast, the diversity of mimetic colour patterns observed in *Heliconius* has arisen from a single “toolkit” of loci that has repeatedly been used to produce both convergent and divergent colour pattern forms within the genus (9). The *optix* gene has recently been identified as one of the these loci, controlling red colour pattern variation (10). Optix is a homeobox transcription factor involved in eye development in *Drosophila* (11), which was co-opted to control scale cell differentiation within the Lepidoptera, and only within *Heliconius* has it taken on a role in colour patterning (12).

The other major colour pattern locus within *Heliconius* (*HeCr/HmYb/N/HnP*) controls a diversity of white and yellow colour pattern elements in the co-mimics *H. erato* (*He*)and *H. melpomene* (*Hm*), but also acts as a supergene controlling polymorphic colour variation in *H. numata* (*Hn*), (13). This locus has more varied effects on colour pattern than *optix*. It controls the presence of the yellow bar on the hind-wing in both *He* (*HeCr*), and *Hm* (*HmYb*), but in *Hm* can also control the presence of a yellow or white forewing band, and even affects the size and distribution of certain red pattern elements (*HmN*) (14,15). In *Hn* it controls black, yellow and orange elements on both wings (*HnP*), producing very different phenotypes that mimic butterflies in the genus *Melinaea* (13). The role of this locus in controlling colour patterning within the Lepidoptera also appears to pre-date other known loci. Genetic variation underlying the *Bigeye* wing pattern mutation in *Bicyclus anynana* and melanism in the peppered moth, *Biston betularia*, both map to homologous genomic regions (16,17) (Figure 1). Therefore this genomic region appears to contain one or more genes that act as major regulators of wing pigmentation and patterning and have done so since early in the evolution of Lepidoptera. This locus has also repeatedly been the target of natural selection, at least three times independently in *Heliconius,* bringing about aposematic colouration and mimicry (18,19), and at least once in a very distantly related moth, maintaining crypsis in varying environments (4). This makes identifying the functional elements of great interest as it will provide further insights into the evolutionary origins of novel traits and the types of genes that act as major targets for natural selection in wild populations.

**Figure 1.**
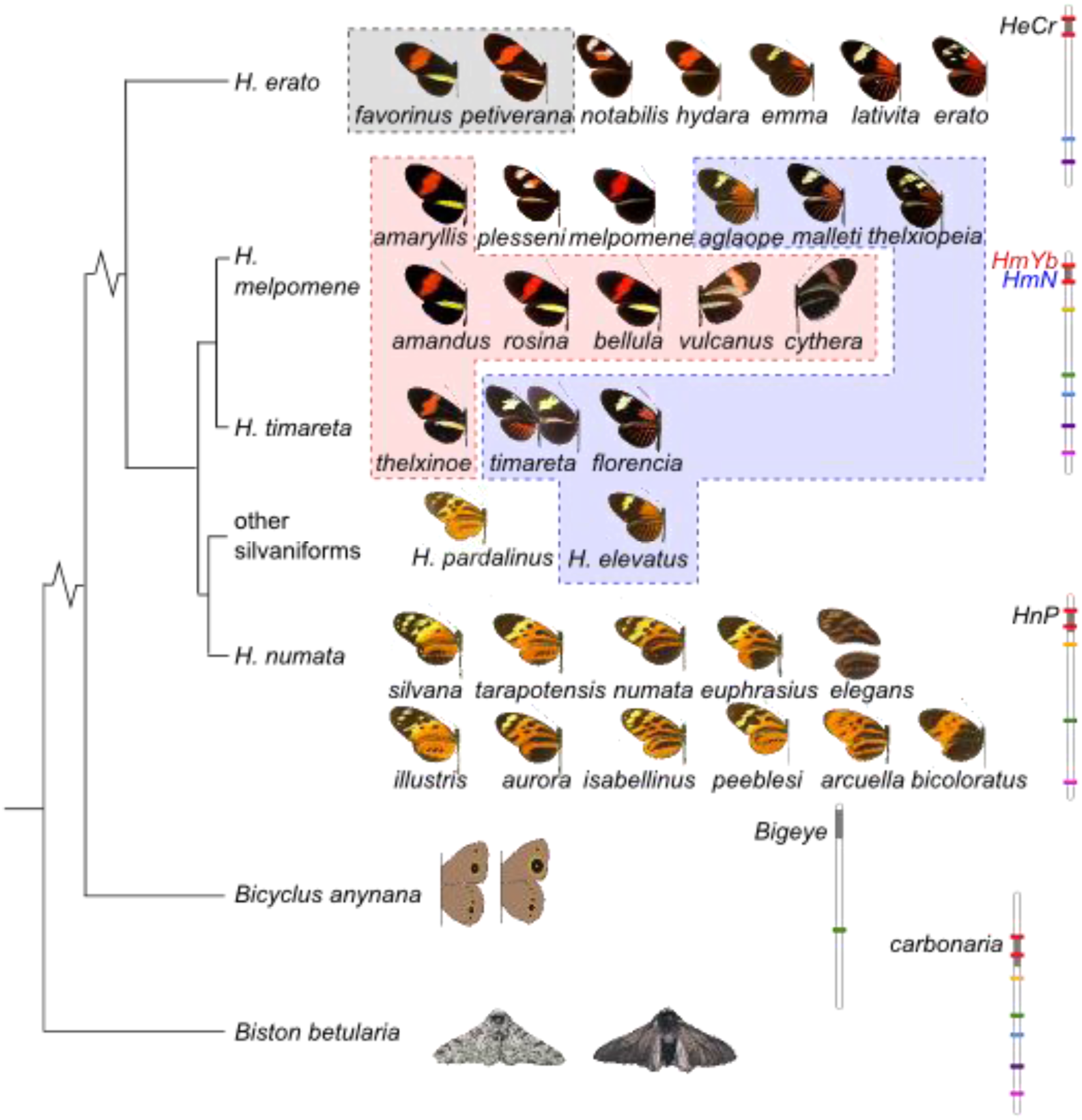
A homologous genomic region controls a diversity of phenotypes across the Lepidoptera. The phylogenetic relationships of the species are shown to the left. On the right: Chromosomal regions containing the colour pattern loci are shown in grey, coloured bars represent markers used to assign homology, with the first and last genes from Figure 2 shown in red (from 13,16,17,56). Dorsal wing surfaces are shown for all *Heliconius* races used, except for *Hm cythera* and *vulcanus*, which have a yellow hind-wing bar only on the ventral surface. In *He* the locus controls the yellow hind-wing bar phenotype (grey boxed races). In *Hm* it controls both the yellow hind-wing bar (pink box) and the yellow forewing band (blue box). In *Hn* It modulates black, yellow and orange elements on both wings, producing phenotypes that mimic butterflies in the genus *Melinaea*.

Mapping of the loci in *He, Hm* and *Hn* has identified an overlapping region of ~1Mb (20–22), which also overlaps with the 1.4Mb region containing the *carbonaria* melanism locus in *B. betularia* (17). Some progress has recently been made in narrowing this region in *Heliconius*. In particular, targeted re-sequencing of the entire mapped colour pattern region using individuals of divergent colour pattern races from either side of a hybrid zone in Peru revealed several narrow peaks of divergence containing several possible candidate genes (23). In addition, we recently demonstrated that parts of this region are shared between the closely related species *Hm* and *H. timareta* and also the more distantly related species *Hm* and *H. elevatus*, resulting in convergence and mimicry between these species (24). Looking for regions that are shared both between hybridising species and across populations that share particular colour pattern elements can therefore provide a way of finding functionally important genomic regions (25,26).

In the current study we used inter and intra-specific comparisons to identify the functional genomic region responsible for colour pattern variation. Associations both with genomic sequence variation and gene expression consistently pointed to a single novel gene, which we have named *poikilomousa* (*poik*), in reference to the tradition of naming *Heliconius* species and races after the Greek Muses and the role of the gene in switching colour patterns.

## Results

### Association analyses

We used the diversity and sharing of colour pattern alleles within *Heliconius* to identify functionally important narrow regions within the previously mapped genomic interval known to contain the *HmYb, HeCr* and *HnP* loci (13,20–22). We looked for single nucleotide polymorphisms (SNPs) that showed associations with colour pattern elements across a diversity of phenotypes. This analysis was performed using three *Heliconius* species groups that are likely to have independently evolved polymorphisms in this genomic region: *He, Hm* and *Hn* (Figure 1).

#### H. erato (He)

We extended the existing BAC sequence tilepath for the *HeCr* region (20). Homologous genes were present in the same order and orientation in *He* and *Hm* (Figure 2B,C). The new contig sequence was used as a reference for alignment of genomic sequences from seven *He* colour pattern races, corresponding to four natural hybrid zones between races (Figure 1, Table S1). We compared individuals from the two races with a yellow hind-wing bar to individuals from the five other colour pattern races, which lack the yellow hind-wing bar. The SNP showing the strongest association with this phenotypic grouping was found just upstream of the coding region of the *poik* gene (Figure 2A), but no SNPs were perfectly associated. We then looked for SNPs showing perfect association with the yellow hindwing bar in either of the two races that have this phenotype. There were 15 SNPs that were homozygous for one allele in *He peitverana* and homozygous for the alternative allele in all races lacking the yellow bar and 108 SNPs showing this fixed pattern for *He favorinus.* There was no overlap in the SNPs identified in these two comparisons, supporting previous suggestions that different *HeCr* alleles are responsible for the convergent phenotypes in these two races (27). The *He petiverana* SNPs were scattered across the region (orange points in Figure 2A), but the *He favorinus* SNPs were all clustered around the *poik* gene (purple points in Figure 2A).

**Figure 2.**
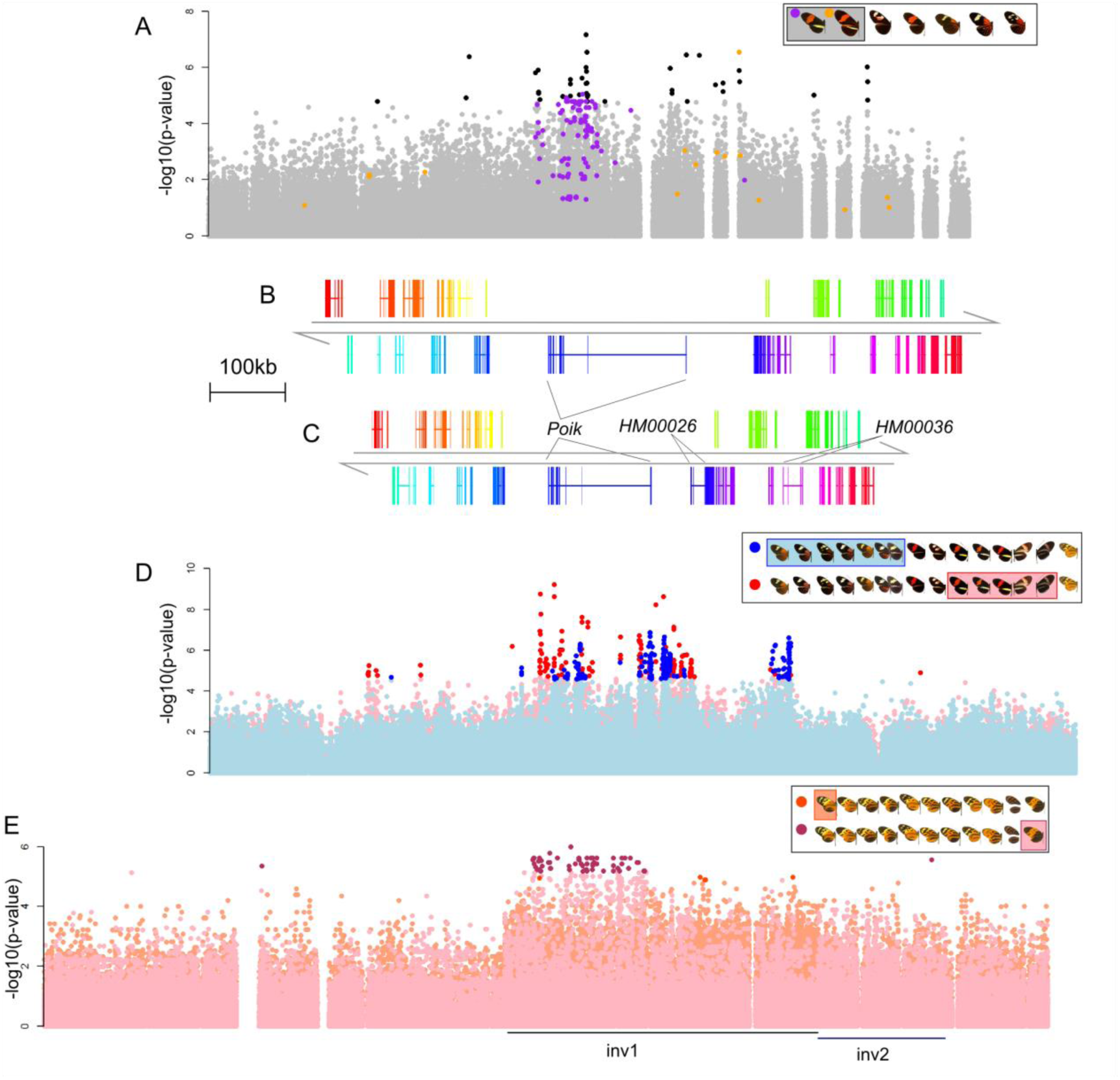
Association analyses across the genomic region known to contain the *HeCr*, *HmYb/Sb/N* and *HnP* loci. A) Association in *He* with the yellow hind-wing bar. Black points are above the level of the strongest association found in non-colour-pattern linked genomic regions. SNPs fixed for a unique state in *He petiverana* are shown in orange and in *He favorinus* in purple. B) Annotation of genes in the *He* BAC walk with direct homologs in *Hm*. Each gene is shown in a different colour with exons (coding and UTRs) connected by a horizontal line, arrows show direction of transcription. C) *Hm* gene positions with colours corresponding to the homologous *He* genes. D) Association in the *Hm/timareta*/silvaniform group with the yellow hind-wing bar (red) and yellow forewing band (blue). Darker coloured points are above the level of the strongest association found in non-colour-pattern linked genomic regions. E) Association with *Hn* morphs. Orange points are associations with the bottom recessive morph (*Hn silvana*) and fuchsia points with the top dominant morph (*Hn bicoloratus*). Positions of the two previously described inversions are shown as black/blue lines below the plot. Darker coloured points are above the level of the strongest association found in non-colour-pattern linked genomic regions.

#### The H. melpomene(Hm)/timareta /silvaniform group

We used a combination of whole-genome and targeted sequencing to obtain sequence data for the *HmYb* region from a diversity of *Hm* races (Figure 1, Table S1). We also included sequences for *H. timareta* and *H. elevatus* (Figure 1), both of which have been shown to exchange colour pattern alleles with *Hm* (24). We tested for genetic associations both with the presence of a yellow hind-wing bar (pink box in Figure 1) and the presence of a yellow forewing band (blue box in Figure 1).

The strongest associations with the yellow hind-wing bar phenotype were found at *poik*, with the most strongly associated SNP found within an intron of this gene (between exons 3 and 4, Figure 2D, Figure 3A). This SNP showed an almost perfect association with the yellow hind-wing bar phenotype except for both *Hm amandus* individuals, which have a yellow bar but were homozygous for the non-yellow bar allele and *H. elevatus,* which was heterozygous (Table S2). Clusters of strongly associated SNPs were also found in the region of the furthest 5’ UTR of this gene (see section on identifying 5’ UTRs) and immediately upstream of this (Figure 2D, red points).

**Figure 3.**
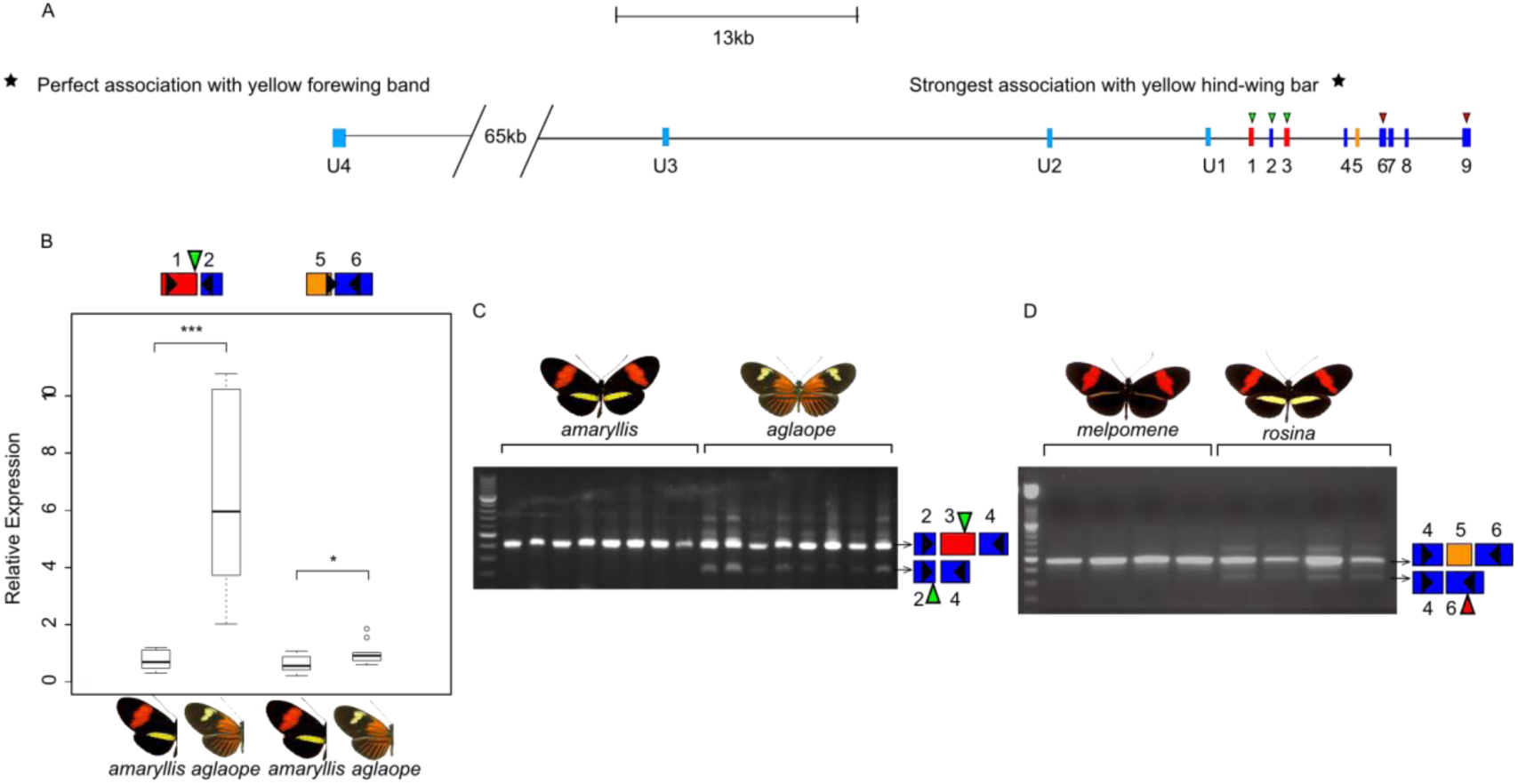
A) Exons and splice variants of *poik* in *Hm*. Orientation is reversed with respect to figures 2 and 4, with transcription going from left to right. SNPs showing the strongest associations with phenotype are shown with stars. B) Differential expression of two regions of *poik* between *Hm amaryllis* and *Hm aglaope* (N=11 and N=10 respectively). C) Expression of a *poik* isoform lacking exon 3 is found in *Hm aglaope* but not *Hm amaryllis*. D) Expression of an isoform lacking exon 5 is found in *Hm rosina* but not *Hm melpomene.* Green triangles indicate predicted start codons and red triangles predicted stop codons, with usage dependent on which exons are present in the isoform. Black triangles indicate the position of the primers used in the assay.

**Figure 4.**
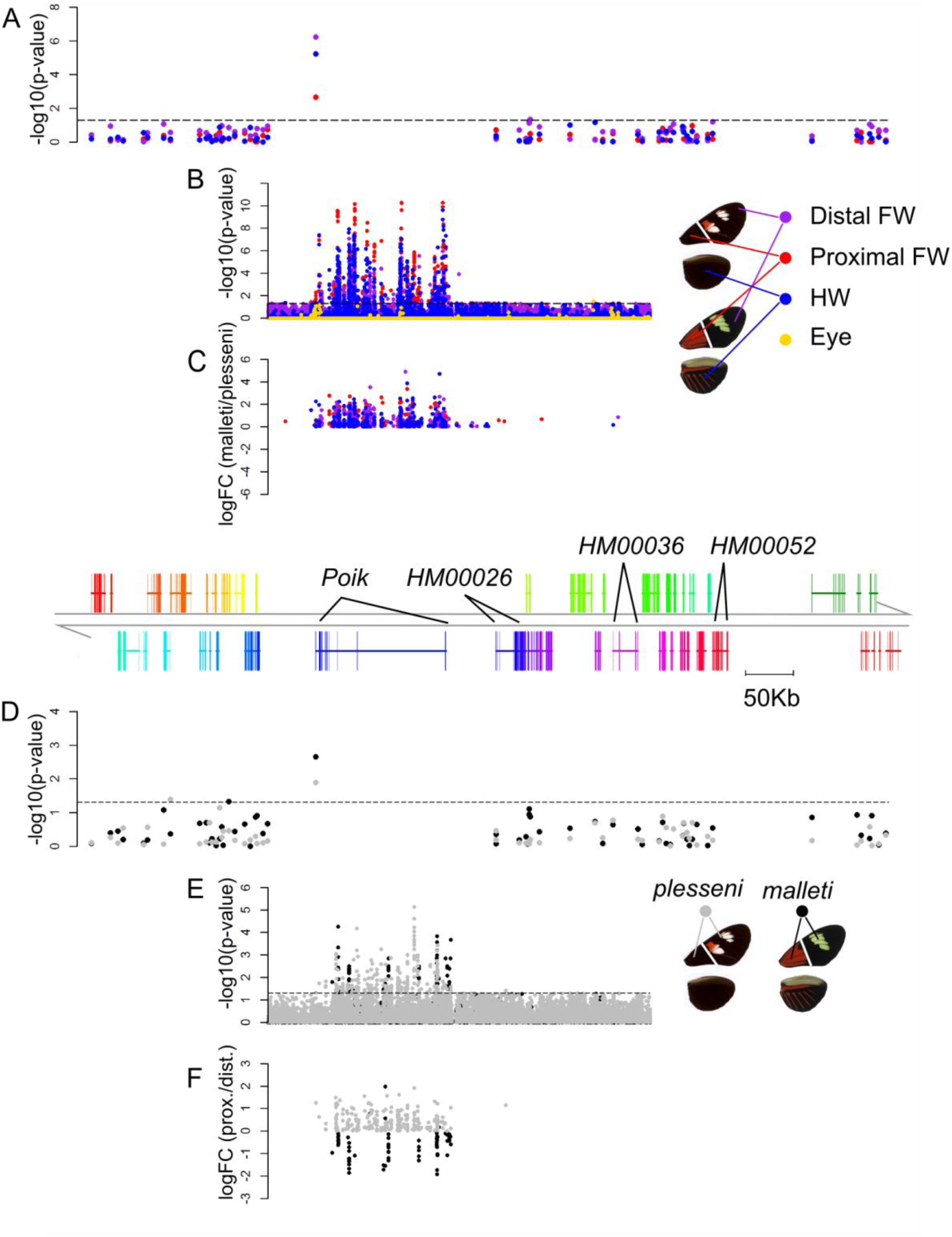
Results for microarray probes from all genes in the *HmYb* interval (A,D) and tiling probes spanning the central portion of the interval (B,C,E,F). Expression is compared between *Hm plesseni* and *Hm malleti* for each wing region (A,B,C) and between proximal and distal forewing regions for each race (D,E,F). Results are from day 3 pupae as these showed the largest differences in expression (see also Figure S2 for other stages). The magnitude and direction of expression difference (log fold-change) is shown for tiling probes showing significant differences (p≤0.05, C,F).

Somewhat similar genomic regions were associated with the presence of the yellow forewing band, in particular the region around the furthest 5’ UTR of *poik.* In addition, strong associations were also found around the more proximal UTR exon U2 (Figure 2D; blue points, Figure 3A). However, strong associations were also found overlapping gene HM00036. A single SNP ~17kb upstream of *poik* (~35kb downstream of HM00026, the next nearest gene) was perfectly associated with the yellow forewing band in all *Hm* races as well as *H. timareta* and *H. elevatus* (Figure 3A, Table S2).

#### H. numata (Hn)

It has previously been shown that large inversions are present at the *HnP* locus between certain *Hn* morphs (22). We found elevated associations with morphs over large genomic blocks corresponding to these inversions (Figure 2E). The top dominant morph, *Hn bicoloratus,* is syntenic with intermediate dominant morphs (eg. *tarapotensis, aurora, acruella*) across inversion 1, and is syntenic with the bottom recessive morphs (eg. *silvana, illustris*) across in version 2 (22). Therefore, it can recombine with all other morphs across one or other of these regions. We observed a distinct narrow region of association with the *bicoloratus* morph, which was above background levels and perfectly corresponded to *poik* (Figure 2E). This associated region does not correspond to any other known genomic feature, such as an inversion or inversion breakpoint.

### 5’ UTRs and alternative splice forms of poik

A previous study of transcriptomic data suggested the existence of different splice variants of *poik* (HM00025) in *Hm* involving both coding exons and alternative 5’ UTR exons (21). We further investigated this using RT-PCR and 5’ RACE on RNA from *Hm* individuals. This revealed an extensive set of alternative 5’ UTRs with the furthest being over 100kb upstream of the *poik* coding exons (Figure 3A). Using the mRNA sequence of these we were able to detect possible homologous regions upstream of the *He poik* gene in the *HeCr* BAC sequence tilepath (Figure 2C), although no corresponding transcripts were found in available RNA-sequencing (RNA-seq) data for *He.*

The furthest upstream exon was present in both *Hm* individuals (*Hm aglaope* and *amaryllis*) used for 5’ RACE and its presence was confirmed by RT-PCR in 17 additional individuals. Moreover exon 1, which contains the start codon, was found to be alternatively spliced with the first UTR exon, in that isoforms contained either exon 1 or exon U1 (Figure 3A). The isoform lacking exon 1 is presumed to utilise the next start codon, which is in exon 3, resulting in a protein that is 365aa rather than 447aa.

We also detected multiple isoforms involving alternative splicing of other coding exons (Figure 3, Figure S1). Isoforms lacking either exon 3 or exon 5 were found to be fairly common and present in multiple individuals. Splicing of exon 3 could lead to a new start codon in exon 2 that would preserve the frame of the rest of the protein and result in a protein of 335aa. Splicing of exon 5 results in a frame shift and premature stop codon in exon 6, and so a truncated protein of 203aa (assuming the exon 1 start codon is used).

### Differential gene expression between H. melpomene races

We conducted independent comparisons of three pairs of *Hm* races that each are found in close geographic proximity and have hybrid zones where genetic exchange occurs (14,19,28). We first used microarrays and RNAseq to investigate expression across the candidate region, with *poik* the only gene to consistently show differences in expression between races. Further comparisons using RT-PCR and qPCR confirmed expression differences at *poik.*

#### Hm plesseni/malleti

These races are from Ecuador and have a hybrid zone on the Eastern slopes of the Andes. They differ at the *HmN* locus, which controls the forewing yellow band and also influences variation in the positioning of red in the forewing band and the length of the hind-wing anterior red bar, and is known to be tightly linked to *HmYb* (14).

We designed a microarray containing probes for all annotated *Hm* genes (24), as well as tiling the central portion of the *HmYb* BAC sequence contig, which was previously identified as showing the strongest differentiation between *Hm* races (23). This was interrogated with RNA from four pupal developmental stages of *Hm plesseni* and *Hm malleti.* We compared levels of gene expression between races for each of three wing regions (hind-wings and two sections of the forewings, Figure 4) and eyes (here used as a non-wing control). *Poik* showed the strongest difference in expression of all genes within the mapped *HmYb/N* interval, with differences between races occurring in all three wing regions at day 1 and day 3 (Figure 4A, Figure S2A). No significant differences in expression of any genes in the region were observed at day 5 or day 7 (Figure S2). This was mirrored in the tiling array results where the strongest differences in expression were found in probes corresponding to *poik,* particularly in the 5’ UTR exons (Figure 4B, Figure S2B). No significant difference in expression was found in the eye. In all cases where differences were detected, *poik* expression was higher in *Hm malleti* than *Hm plesseni* (Figure 4C, Figure S2C). Expression differences within the presumed *poik* introns suggest the presence of additional 5’ UTR exons that were not detected by RACE or RT-PCR.

We also used this data to investigate spatial patterns of gene expression on the wing by comparing gene expression between proximal and distal forewing sections within each race. *Hm malleti* has a forewing yellow bar, controlled by the *HmN* locus, while in *Hm plesseni* this locus controls the positioning of red and white scales within the forewing band region. When comparing expression levels between wing sections across the *HmYb/N* genomic region, significant differences were again found primarily in day 1 and day 3 pupal wings rather than day 5 or day 7 (Figure 4, Figure S2), consistent with the comparisons of gene expression between races, and suggesting that these earlier stages are the most important for pattern specification. Furthermore, *poik* again showed the largest and most significant differences in expression in the *HmYb* region from both the gene array and the tiling array (Figure 4D,E, Figure S2H,I). *Poik* expression in *Hm malleti* was generally higher in the distal section that contains the yellow forewing band, although a few probes showed the opposite pattern, perhaps suggesting wing region specific splicing variation. In contrast for *Hm plesseni* expression was consistently higher in the proximal wing region (Figure 4F, Figure S2J). There was also evidence for differential splicing of *poik* between races, as the regions of *poik* showing differential expression were different between the two races.

#### Hm amaryllis/aglaope

These races have a hybrid zone in Peru and differ at the *HmYb* and *HmN* loci controlling the presence of the yellow hind-wing bar and yellow forewing band respectively. RNA-seq data for hind-wings from three developmental stages had previously been obtained for two individuals of each race at each stage (12 individuals in total) and used in the annotation of the *Hm* genome (24). We analysed this data to look for differences in gene expression between races and detected twelve, 95 and 208 genes as being differentially expressed between races at final instar larvae, day 2 and day 3 respectively using multiple analysis methods (Table S3). Only two genes were detected as being differentially expressed within the *HmYb* mapped region and both were only differentially expressed in the day 2 wings. HM00052 was upregulated in the yellow barred hind-wings of *Hm amaryllis* (p=0.018) while *poik* was upregulated in the rayed hind-wings of *Hm aglaope* (p=0.035). This difference in expression of *poik* is consistent with the upregulation that we detected in the phenotypically similar *Hm malleti*, and could be linked to the role of the *HmYb/N* locus in controlling the length of the hind-wing anterior red bar (14).

The *poik* expression difference was confirmed by quantitative RT-PCR (qPCR) using day 2 hind-wings from 10 *Hm aglaope* and 11 *Hm amaryllis*. On average expression was 1.6 times higher in *Hm aglaope* (SD=0.7, Wilcoxon rank sum test p=0.035) using primers in the coding exons 5 and 6 (Figure 3B). However, using the same samples, we found 8.5x higher expression in *Hm aglaope* when assaying exons 1 and 2 (SD=0.54, Wilcoxon rank sum test p=1.08e-05, Figure 3B). This suggests that *Hm aglaope* and *Hm amaryllis* have differential expression of the isoforms that contain alternative exons 1 and U1, which contain different start codons.

In addition we found that the isoform lacking exon 3 was differentially expressed between these races. It was detected in all rayed *Hm aglaope* individuals (developing hind-wings from final instar larvae, day 1 and day 2 pupae, 24 individuals in total) but appeared to be completely absent from all yellow barred *Hm amaryllis* (same stages and sample sizes used, Figure 3C, Figure S1B).

#### Hm rosina/melpomene

These races have a hybrid zone in Panama and differ only in the presence of the yellow hind-wing bar, with *Hm rosina* having a bar and *Hm melpomene* lacking it. Comparisons of these races were conducted by RT-PCR and qPCR of *poik* transcripts only. Unlike the previous comparison no difference in expression was detected when using assays spanning either exons 5 and 6 or exons 1 and 2 (Day 2 pupal wings, n=25, Wilcoxon rank sum test p=0.8517 and p=0.205 respectively). Neither was there any clear race association with the isoform lacking exon 3, with a limited number of both *Hm melpomene* and *rosina* expressing this isoform (Figure S1C). This could suggest that these differences that were detected in the previous comparisons are associated with the control of the shape of the anterior red bar on the hind-wing that is present in both *Hm aglaope* and *malleti* but not in either *Hm rosina* nor *melpomene.*

However, in this comparison we did detect one isoform that was differentially expressed between races. An isoform lacking exon 5 was detected in all *Hm rosina* individuals, which have a yellow hind-wing bar (developing hind-wings from final instar larvae, day 1 and day 2 pupae, 17 individuals in total) but was not present in any *Hm melpomene* individuals, which lack the bar (same stages and sample size). This isoform showed allele specific expression in an F2 cross between *Hm rosina* and *Hm melpomene,* demonstrating *cis*-regulatory control of the alternative splicing patterns. Using markers within the *HmYb* region we were able to identify individuals as heterozygous or homozygous for *HmYb* from the parental populations. Individuals both hetero- and homozygous for the *Hm rosina* allele expressed the isoform lacking exon 5, while those homozygous for the *Hm melpomene* allele did not (Figure S1H). Using a diagnostic SNP within exon 4, we found that in heterozygous individuals only the *Hm rosina* allele produced this isoform, while other isoforms contained alleles from both parents (Figure S1I).

We also found the isoform lacking exon 5 to be expressed in *Hm cythera* (pool of 17, and 2 further individuals), which again possess the yellow hind-wing bar, and to be absent from a pool of 6 *Hm malleti* individuals, which lack the bar (Figure S1G). However, we did not find a consistent difference in expression of this isoform between *Hm aglaope* and *amaryllis* (Figure S1F), although the lower expression detected at exons 5 and 6 in *Hm amaryllis* (Figure 3B) could indicate relatively higher prevalence of isoforms lacking exon 5 in this race. Therefore, isoforms lacking exon 5 may be important in formation of the yellow hind-wing bar.

## Discussion

### Identification of a gene involved in lepidopteran scale pigmentation and patterning

We have identified a gene that we name *poik,* which underlies pigmentation patterning in *Heliconius.* Focusing on a region previously shown to control colour pattern variation in *Heliconius* (13,21), we find consistent associations between DNA sequence variation at *poik* and a diversity of colour pattern variation across multiple species. This includes associations with the presence of the yellow hind-wing bar in both of the co-mimics *He* and *Hm* and also with the presence of the yellow forewing band only in *Hm (He* is known to control this bar using alternative loci (29)). In addition, in *Hn* we find strong morph associations specifically highlighting *poik* within the larger, previously identified inversions in this region (22). We also find differences in expression of *poik* associated with colour pattern variation. In three pairwise comparisons of closely related *Hm* races, *poik* is the only gene in this region to show consistent differential expression. In addition, we find differential expression of *poik* across the developing wing in a pattern corresponding to adult colour pattern elements, strengthening the inference that *poik* is involved in specifying colour pattern.

Most previously identified wing patterning genes have been transcription factors or signalling molecules. In contrast, the closest orthologues of *poik* are cell cycle regulatory proteins including the *Drosophila* gene *cortex* and the fizzy family, making this a surprising candidate for controlling wing patterning.

### A novel function for a member of a conserved cell cycle regulator family

We explored the origin of the *poik* gene and show that it falls in an insect specific lineage within the fizzy family of cell cycle regulators (Figure 5). The phylogenetic tree of the gene family highlighted three major orthologous groups. Two of these represent highly conserved proteins, one containing human and yeast CDC20 and *Drosophila* fzy, the other containing orthologues of cdh1/fzr/rap. CDC20/fzy has a highly conserved function in cell cycle regulation, which involves targeting specific proteins, including cyclins and other cell stage specific proteins, for degradation. This is mediated through interaction with the anaphase promoting complex/cyclosome (APC/C) and acts to regulate exit from mitosis (30,31).

**Figure 5.**
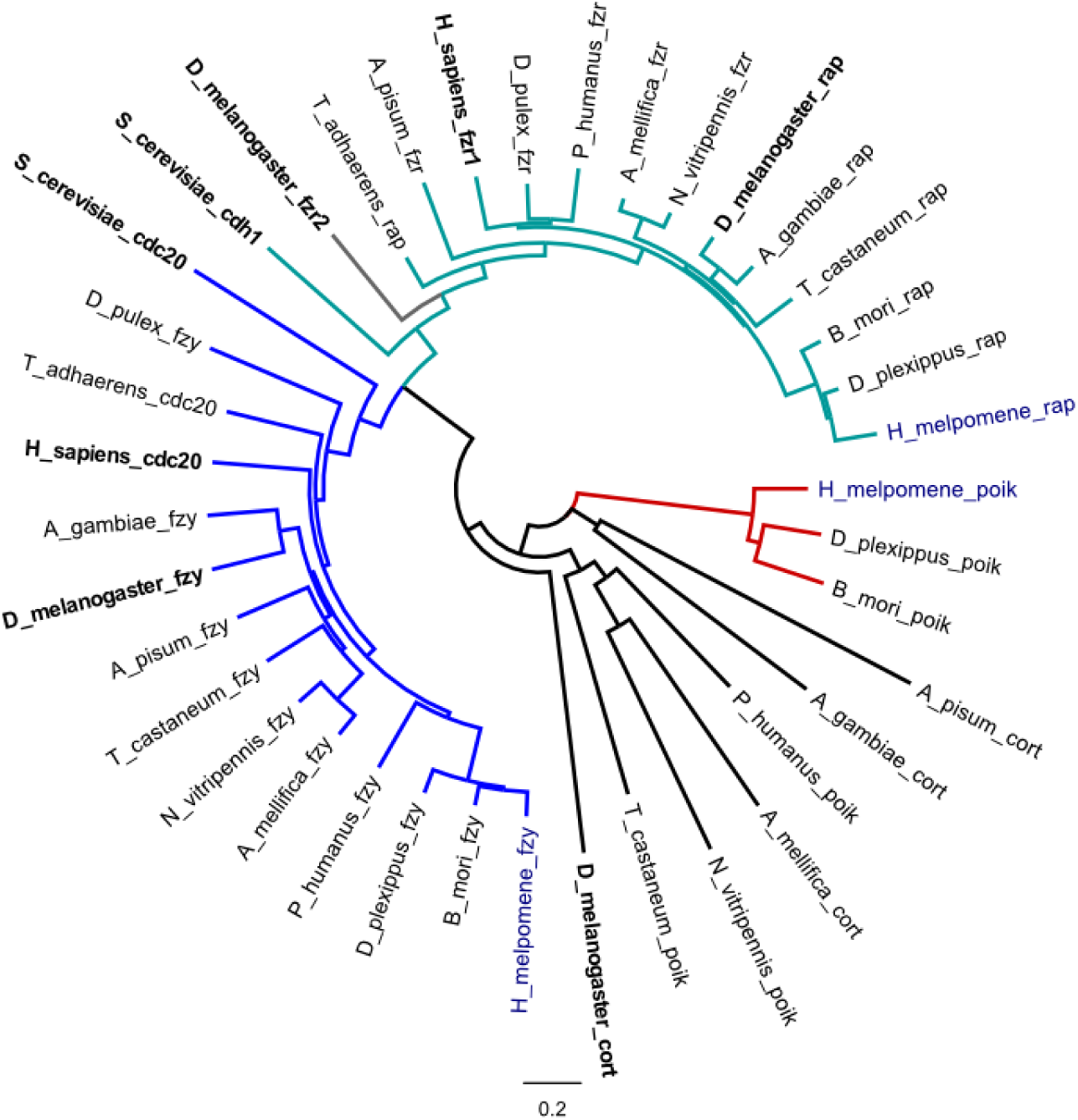
Neighbour joining phylogeny of Fizzy family proteins including functionally characterised proteins (in bold) from *Saccharomyces cerevisiae, Homo sapiens* and *Drosophila melanogaster* as well as copies from the basal metazoan *Trichoplax adhaerens* and a range of annotated arthropod genomes (*Daphnia pulex, Acyrthosiphon pisum, Pediculus humanus, Apis mellifica, Nasonia vitripennis, Anopheles gambiae, Tribolium castaneum*) including the lepidoptera *H. melpomene* (in blue), *Danaus plexippus* and *Bombyx mori.* Branch colours: dark blue, CDC20/fzy; light blue, CDH1/fzr/rap; red, lepidoptran poik. Made using Geneious v6.1 (Biomatters).

Cdh1/fzr/rap are also highly conserved proteins found across eukaryotes (Figure 5) and are very similar in function to CDC20/fzy but appear to be slightly less conserved in their targets and may have tissue specific effects (32–34). Conserved orthologues of both of these were found in the *Hm* and other lepidopteran genomes (on *Hm* chromosomes 11 and 18, see Extended Experimental Procedures). Poik appears to belong to a third group, which was only identified in the insect genomes. The only functionally characterised member of this group is the *Cortex* gene (*Cort*) in *Drosophila melanogaster.* This acts through a similar mechanism to CDC20/fzy in order to control meiosis in the female germ line (35–37). *D. melanogaster* Cort falls as a highly divergent outgroup to the other predicted insect proteins in this group. Indeed, *Hm* poik shows similarly low amino acid sequence identity to *D. melanogaster* Cort (11.6%) and *D. melanogaster* fzy (14.4%) (Figure 5).

Overall, the phylogenetic patterns suggest that poik is a distantly related member of the fizzy family of proteins, belonging to a group that is unique to the insects. These have faster evolutionary rates than other members of this family, with the low amino acid identity between *D. melanogaster* cort and *H. melpomene* poik (11.6%) contrasting with much higher identities for fzy (46.7%) and rap/fzr (47.5%) (Figure 5, Figure S3). Fast evolutionary rates for *poik* have also been found previously (38). However, poik does have some conservation of the fizzy family C-box and IR elements that mediate binding to the APC/C (36), suggesting that it may have retained the ability to bind to this complex (Figure S3).

It is conceivable that within the lepidoptra, *poik* may have been co-opted to control scale cell development. Wing scales have fairly peculiar development as they arise from greatly expanded cells that become highly polyploid during development (1). The enlarged polyploid cells are similar to the phenotype observed when *Cort* or *fzr* are overexpressed in *D. melanogaster* wings or other tissues (34,37). We found that expressing *Hm poik* in *D. melanogaster* wings produced no phenotypic effect (Extended Experimental Procedures, Figure S4), but this may simply be due to lack of conservation of its binding partners or targets.

We propose that *poik* controls pigmentation patterning on the wing through regulation of cell cycle timing in developing scale cells. Scales of different colours develop at different rates in both butterflies and moths, with pale coloured scales developing earlier than melanic scales in all Lepidoptera studied so far, including *Heliconius* (1). The timing of differential expression of *poik* in early pupal development (up to day 3) is consistent with a role in controlling the timing of scale development (2). Regulation of scale development rate has previously been proposed as a mechanism for control of colour patterning (39,40). Therefore, it seems likely that regulatory changes that alter either the timing, expression level or splicing of *poik* in a pattern specific way could bring about differences in cell development rate and so alter colour pattern. The precise mechanism remains unknown, but could either act during the two cell divisions involved in scale cell development, or play a role in the timing of wing scale maturation, which differs between wing regions (Aymone et al., 2013). There is a precedent for involvement of this gene family in developmental patterning, as *rap/fzr* controls pattern formation in the *D. melanogaster* developing eye-antennal disk (34).

### Alternative splicing as a target of selection and a means of generating diversity

Alternative splicing has often been proposed as a mechanism for generating additional diversity from the same basic genetic toolkit and so a potential target for natural selection (41). In the swallowtail butterflies, splicing differences are involved in polymorphic colour pattern mimicry (6). In *Heliconius* we have found high levels of alternative splicing of *poik,* with some isoforms showing race and possible wing region specific expression, suggesting that splicing of this gene may also be important in generating adaptive wing pattern diversity. Splicing patterns are known to be strongly influenced by intronic sequence variation (41), so the presence of phenotypically associated SNPs in introns of *poik* could regulate differences in splicing and play a role in generating differences between races.

The race-associated isoforms that we observed all resulted in altered protein sequences. Fizzy family proteins contain multiple dispersed domains that mediate the interactions with proteins that are targeted for degradation (30,42–44), so different isoforms could contain different combinations of these domains, possibly altering their specificity for particular targets. It has also been shown that modified fizzy family proteins that lack the domains for binding to the APC/C, can instead reduce degradation of target proteins by blocking the binding sites of these proteins (43). The truncated isoform found in *H. m. rosina* lacks the IR tail, which mediates binding to the APC/C (36), so may have such a function.

### Architecture of a “supergene”

Supergenes are single Mendelian loci that control complex alternative adaptive phenotypes (45), and were initially proposed to be assemblages of several tightly linked genes acting together to control switches between complex phenotypes (46). The inversions present between *Hn* morphs supported this model by providing a mechanism whereby multiple genes could be held together in tight linkage (22). However our findings support those from *Papilio* suggesting that a supergene may in fact be a single gene with multiple downstream targets that enable it to control a complex phenotypic polymorphism (6). The alternative *poik* 5’ UTRs that we detected suggest multiple regulatory regions spread over more than 100Kb. These likely contain binding sites for multiple upstream regulators, facilitating the production of a diversity of downstream patterns. Under this model the inversions present in *Hn* could be acting to couple together the large 5’ regulatory region of *poik.*

Nevertheless it remains possible that there are also additional functional genes controlling patterning in this region in both *Hn* and in species with no supergene architecture. For example, in *Hm* we find strong phenotypic associations around gene *HM00036* as well as *poik.* Nonetheless, *poik* is the only gene for which we detect DNA sequence variation consistently associated with colour pattern in multiple comparisons, and also expression differences associated with both race and wing region.

### Conclusions

We have identified a gene, *poik,* controlling colour pattern variation in *Heliconius.* This gene is a member of a conserved cell cycle regulator family and is most likely an orthologue of a female germ line specific protein controlling the switch from mitosis to meiosis in *Drosophila.* Therefore this gene was not a likely *a priori* candidate, suggesting that evolution can be unpredictable in its targets, with unexpected genes taking on novel functions. Nevertheless, following a possible switch to a role in scale development, *poik* has repeatedly been targeted by natural selection acting on colour pattern. Previous work showing broad homology between this locus in *Heliconius* and the *carbonaria* locus in the peppered moth (17) suggests that *poik* may act as a major switch in wing pattern evolution across the Lepidoptera.

## Materials and Methods

Detailed protocols and procedures are included in Extended Experimental Procedures in the Supplemental Information.

### Association analyses

We measured associations between genotype and phenotype using a score test (qtscore) in the GenABEL package in R (47). This was corrected for background population structure using a test specific inflation factor, *λ*, calculated from genomic regions unlinked to the major colour pattern controlling loci, as the colour pattern loci are known to have different population structure to the rest of the genome (24–26). Information on the individuals used and ENA accessions for sequence data are given in Table S1.

#### He

We used an existing *He* BAC library (20) to identify BACs extending the sequenced *HeCr* region, which were then sequenced and assembled. One gap remained in the reference (between positions 800,387 and 848,446), which was filled using scaffolds from an initial assembly of the *He* genome. Homology and synteny with the *Hm* reference were identified by aligning the *Hm* coding sequences to the *He* reference with BLAST. The Cr contig is deposited in Genbank with accession KC469893.

We used shotgun Illumina sequence reads from 45 *He* individuals from 7 races that were generated as part of a previous study (48)(Table S1). Reads were aligned to an *He* reference containing the *HeCr* contig and other sequenced *He* BACs (20,48).

#### Hm/timareta/silvaniform clade

We used previously published sequence data from targeted sequencing of the *HmYb/Sb/N* and *HmB/D* colour pattern loci and ~1.8Mb of non-colour pattern genomic regions (23), as well as whole genome shotgun sequencing (24,49). We also added further targeted sequencing and shotgun whole genome sequencing of additional individuals (Table S1). Reads were aligned to either v1.1 of the *Hm* reference genome, in the case of *Hn,* or to a reference genome with the scaffolds containing *HmYb/N* and *HmB/D* swapped with reference BAC sequences (24), in the case of *Hm/timareta*/silvaniform. We used the BAC sequences of the colour pattern interval for the *Hm/ timareta* /silvaniform analysis because this contains fewer gaps of unknown sequence. However, we used the genome scaffold for the *Hn* analysis because this is longer making it easier to compare the inverted and non-inverted regions present in this species. A total of 49 individuals were included in the *HmYb/N* association analysis of the *Hm/timareta* /silvaniform clade and a total of 26 *Hn* individuals were included in the analysis of the *HnP* locus (Figure 1 and Table S1). We tested for associations with the *Hn* two morphs with the largest sample sizes (*Hn silvana,* n=4 and *Hn bicoloratus,* n=5)

### Gene Expression Analyses

All tissues used for gene expression analyses were dissected from individuals from captive stocks derived from wild caught individuals of the various races of *Hm* (*aglaope, amaryllis, melpomene, rosina, plesseni, malleti*).

#### Tiling microarray

Samples were labelled with Cy3 and each hybridised to a separate array. The *HmYb* probe array contained 9,979 probes distanced on average at 10bp. The whole-genome expression array contained on average 9 probes per annotated gene in the genome (v1.1 (24)) as well as any transcripts not annotated but predicted from RNA-seq evidence. The microarray data are deposited in GEO with accessions GSM1563402-GSM1563497.

The tiling array and whole-genome data sets were analysed separately. Expression values were extracted and quantile-normalised, log_2_-transformed, quality controlled and analysed for differences in expression between individuals and wing regions. P-values were adjusted for multiple hypotheses testing using the False Discovery Rate (FDR) method (50).

#### RNA-sequencing

RNAseq reads are deposited in ENA under study accessions ERP000993 and PRJEB7951. Two methods were used for alignment of reads to the reference genome and inferring read counts, Stampy (51) and RSEM (RNAseq by Expectation Maximisation) (52). In addition we used two different R/Bioconductor packages for estimation of differential gene expression, DESeq (53) and BaySeq (54). We present results for the genes detected as differentially expressed with all four methods (see Table S3 for results from each method).

#### 5’ RACE, RT-PCR and qPCR

Total RNA was extracted from hindwings from captive stocks including F2 individuals from a *Hm rosina* (female) x *Hm melpomene* (male) cross. RNA was thoroughly checked for DNA contamination before synthesising single stranded cDNA with random (N6) primers. cDNA was used for RT-PCR and qPCR using gene (or isoform) specific primers (Table S4). For qPCR we used two housekeeping genes (*EF1α* and *Ribosomal Protein S3A*) for normalisation and all results were taken as averages of triplicate PCR reactions for each sample. Statistical significance was assessed by Wilcoxon rank sum tests performed in R (55).

5’ RACE was performed using RNA from hind-wing discs from one *Hm aglaope* and one *Hm amaryllis* final instar larvae. We identified isoforms from 5’ RACE and RT-PCR products by cutting individual bands from agarose gels and if necessary by cloning products before Sanger sequencing. Sequenced isoforms are deposited in GenBank with accessions XXXXXXXX. The presence of the furthest 5’ UTR exon was confirmed in 17 individuals comprising *Hm aglaope* and *Hm amaryllis* of various developmental stages.

## Acknowledgements

We thank Christopher Saski, Clemson University, for assembly of the *He* BACs. Richard Merrill, Moises Abanto and Adriana Tapia assisted with raising butterflies. Anna Morrison, Robert Tetley and Sarah Carl assisted with lab work at the University of Cambridge. We thank the governments of Colombia, Ecuador, Panama and Peru for permission to collect butterflies. This work was funded by a Leverhulme Trust award and BBSRC grant (H01439X/1) to CDJ, NSF grants (DEB 1257689, IOS 1052541) to WOM and an ERC starting grant to MJ. NJN is funded by a NERC fellowship (NE/K008498/1).

## Abbreviations

*He, Heliconius erato*

*Hm, Heliconius melpomene*

*Hn, Heliconius numata*

*poik, poikilomousa*

## Supporting Information

**SI Supporting information: contains Extended Experimental Procedures; Supplemetal Figures S1, S2 and S4; Supplemental tables S2, S3 and S4**

**Table S1. Samples used for association analyses**

**Figure S3. Fizzy family protein alignments.** Annotations with names underlined in red were manually adjusted based on BLAST search results using functionally characterised proteins. The C-box region is outlined in red (DR(F/Y)IPXRX[~45-75a.a.](K/R)XL), and the IR tail in blue, which mediate binding to the APC/C. Alignment made using Geneious v6.1 (Biomatters).

## References

1. Nijhout HF. The development and evolution of butterfly wing patterns. Washington: Smithsonian Institution Press; 1991.

2. Aymone ACB, Valente VLS, de Araújo AM. Ultrastructure and morphogenesis of the wing scales in Heliconius erato phyllis (Lepidoptera: Nymphalidae): What silvery/brownish surfaces can tell us about the development of color patterning? Arthropod Struct Dev. 2013;42(5):349–359.

3. Olofsson M, Løvlie H, Tibblin J, Jakobsson S, Wiklund C. Eyespot display in the peacock butterfly triggers antipredator behaviors in naïve adult fowl. Behav Ecol. 2013;24(1):305–310.

4. Cook LM, Grant BS, Saccheri IJ, Mallet J. Selective bird predation on the peppered moth: the last experiment of Michael Majerus. Biol Lett. 2012;8(4):609–612.

5. Jiggins CD. Ecological Speciation in Mimetic Butterflies. BioScience. 2008;58(6):541–548.

6. Kunte K, Zhang W, Tenger-Trolander A, Palmer DH, Martin A, Reed RD, et al. doublesex is a mimicry supergene. Nature. 2014;507(7491):229–232.

7. Clark R, Brown SM, Collins SC, Jiggins CD, Heckel DG, Vogler AP. Colour pattern specification in the Mocker swallowtail Papilio dardanus: the transcription factor invected is a candidate for the mimicry locus H. Proc R Soc B Biol Sci. 2008;275(1639):1181–1188.

8. Scriber JM, Hagen RH, Lederhouse RC. Genetics of Mimicry in the Tiger Swallowtail Butterflies, Papilio glaucus and P. canadensis (Lepidoptera: Papilionidae). Evolution. 1996;50(1):222–236.

9. Joron M, Jiggins CD, Papanicolaou A, McMillan WO. Heliconius wing patterns: an evo-devo model for understanding phenotypic diversity. Heredity. 2006;97(3):157–167.

10. Reed RD, Papa R, Martin A, Hines HM, Counterman BA, Pardo-Diaz C, et al. optix Drives the Repeated Convergent Evolution of Butterfly Wing Pattern Mimicry. Science. 2011;333(6046):1137–1141.

11. Seimiya M, Gehring WJ. The Drosophila homeobox gene optix is capable of inducing ectopic eyes by an eyeless-independent mechanism. Development. 2000;127(9):1879–1886.

12. Martin A, McCulloch KJ, Patel NH, Briscoe AD, Gilbert LE, Reed RD. Multiple recent co-options of Optix associated with novel traits in adaptive butterfly wing radiations. EvoDevo. 2014;5(1):7.

13. Joron M, Papa R, Beltrán M, Chamberlain N, Mavárez J, Baxter S, et al. A Conserved Supergene Locus Controls Colour Pattern Diversity in *Heliconius* Butterflies. PLoS Biol. 2006;4(10). doi: 10.1371/journal.pbio.0040303

14. Nadeau NJ, Ruiz M, Salazar P, Counterman B, Medina JA, Ortiz-Zuazaga H, et al. Population genomics of parallel hybrid zones in the mimetic butterflies, *H. melpomene* and *H. erato*. Genome Res. 2014;24(8):1316–1333.

15. Sheppard PM, Turner JRG, Brown KS, Benson WW, Singer MC. Genetics and the Evolution of Muellerian Mimicry in *Heliconius* Butterflies. Philos Trans R Soc Lond B Biol Sci. 1985;308(1137):433–610.

16. Beldade P, Saenko SV, Pul N, Long AD. A Gene-Based Linkage Map for *Bicyclus anynana* Butterflies Allows for a Comprehensive Analysis of Synteny with the Lepidopteran Reference Genome. PLoS Genet. 2009;5(2):e1000366. doi: 10.1371/journal.pgen.1000366

17. Van’t Hof AE, Edmonds N, Dalíková M, Marec F, Saccheri IJ. Industrial Melanism in British Peppered Moths Has a Singular and Recent Mutational Origin. Science. 2011 20;332(6032):958–60.

18. Kapan DD. Three-butterfly system provides a field test of mullerian mimicry. Nature. 2001;409(6818):338–340.

19. Mallet J, Barton NH. Strong Natural Selection in a Warning-Color Hybrid Zone. Evolution. 1989;43(2):421–431.

20. Counterman BA, Araujo-Perez F, Hines HM, Baxter SW, Morrison CM, Lindstrom DP, et al. Genomic Hotspots for Adaptation: The Population Genetics of Müllerian Mimicry in *Heliconius erato*. PLoS Genet. 2010;6(2):e1000796. doi: 10.1371/journal.pgen.1000796

21. Ferguson L, Lee SF, Chamberlain N, Nadeau NJ, Joron M, Baxter S, et al. Characterization of a hotspot for mimicry: assembly of a butterfly wing transcriptome to genomic sequence at the HmYb/Sb locus. Mol Ecol. 2010;19(s1):240–254. doi: 10.1111/j.1365-294X.2009.04475.x

22. Joron M, Frezal L, Jones RT, Chamberlain NL, Lee SF, Haag CR, et al. Chromosomal rearrangements maintain a polymorphic supergene controlling butterfly mimicry. Nature. 2011; 477 (7363):203–206.

23. Nadeau NJ, Whibley A, Jones RT, Davey JW, Dasmahapatra KK, Baxter SW, et al. Genomic islands of divergence in hybridizing Heliconius butterflies identified by large-scale targeted sequencing. Philos Trans R Soc B Biol Sci. 2012;367(1587):343–353.

24. Consortium THG. Butterfly genome reveals promiscuous exchange of mimicry adaptations among species. Nature. 2012;487(7405):94–98.

25. Hines HM, Counterman BA, Papa R, Albuquerque de Moura P, Cardoso MZ, Linares M, et al. Wing patterning gene redefines the mimetic history of Heliconius butterflies. Proc Natl Acad Sci. 2011;108(49):19666–19671.

26. Pardo-Diaz C, Salazar C, Baxter SW, Merot C, Figueiredo-Ready W, Joron M, et al. Adaptive Introgression across Species Boundaries in Heliconius Butterflies. PLoS Genet. 2012;8(6):e1002752. doi: 10.1371/journal.pgen.1002752

27. Maroja LS, Alschuler R, McMillan WO, Jiggins CD. Partial Complementarity of the Mimetic Yellow Bar Phenotype in Heliconius Butterflies. PLoS ONE. 2012;7(10):e48627. doi: 10.1111/j.1558-5646.2009.00767.x

28. Mallet J. Hybrid zones of *Heliconius* butterflies in Panama and the stability and movement of warning colour clines. Heredity. 1986;56(2):191–202.

29. Mallet J. The Genetics of Warning Colour in Peruvian Hybrid Zones of Heliconius erato and H. melpomene. Proc R Soc Lond B Biol Sci. 1989;236(1283):163–185.

30. Barford D. Structural insights into anaphase-promoting complex function and mechanism. Philos Trans R Soc B Biol Sci. 2011;366(1584):3605–3624.

31. Dawson IA, Roth S, Artavanis-Tsakonas S. The Drosophila Cell Cycle Gene fizzy Is Required for Normal Degradation of Cyclins A and B during Mitosis and Has Homology to the *CDC20* Gene of *Saccharomyces cerevisiae*. J Cell Biol. 1995;129(3):725–737.

32. Jacobs H, Richter D, Venkatesh T, Lehner C. Completion of mitosis requires neither fzr/rap nor fzr2, a male germline-specific Drosophila Cdh1 homolog. Curr Biol CB. 2002;12(16):1435–1441.

33. Li R, Wan B, Zhou J, Wang Y, Luo T, Gu X, et al. APC/C(Cdh1) targets brain-specific kinase 2 (BRSK2) for degradation via the ubiquitin-proteasome pathway. PloS One. 2012;7(9):e45932. doi: 10.1371/journal.pone.0045932

34. Pimentel AC, Venkatesh TR. rap gene encodes Fizzy-related protein (Fzr) and regulates cell proliferation and pattern formation in the developing Drosophila eye-antennal disc. Dev Biol. 2005; 285(2):436–446.

35. Chu T, Henrion G, Haegeli V, Strickland S. Cortex, a Drosophila gene required to complete oocyte meiosis, is a member of the Cdc20/fizzy protein family. Genesis. 2001;29(3):141–152.

36. Pesin JA, Orr-Weaver TL. Developmental Role and Regulation of cortex, a Meiosis-Specific Anaphase-Promoting Complex/Cyclosome Activator. PLoS Genet. 2007;3(11):e202. doi: 10.1371/journal.pgen.0030202

37. Swan A, Schüpbach T. The Cdc20/Cdh1-related protein, Cort, cooperates with Cdc20/Fzy in cyclin destruction and anaphase progression in meiosis I and II in Drosophila. Development 2007;134(5):891–899.

38. Wu G, Joron M, Jiggins C. Signatures of selection in loci governing major colour patterns in Heliconius butterflies and related species. BMC Evol Biol. 2010;10(1):368.

39. Gilbert LE, Forrest HS, Schultz TD, Harvey DJ. Correlations of ultrastructure and pigmentation suggest how genes control development of wing scales of Heliconius butterflies. J Res Lepidoptera. 1988;26:141–160.

40. Koch PB, Lorenz U, Brakefield PM, ffrench-Constant RH. Butterfly wing pattern mutants: developmental heterochrony and co-ordinately regulated phenotypes. Dev Genes Evol. 2000;210(11):536–544.

41. Keren H, Lev-Maor G, Ast G. Alternative splicing and evolution: diversification, exon definition and function. Nat Rev Genet. 2010;11(5):345–355.

42. Kraft C, Vodermaier HC, Maurer-Stroh S, Eisenhaber F, Peters J-M. The WD40 Propeller Domain of Cdh1 Functions as a Destruction Box Receptor for APC/C Substrates. Mol Cell. 2005;18(5):543–553.

43. Pfleger CM, Lee E, Kirschner MW. Substrate recognition by the Cdc20 and Cdh1 components of the anaphase-promoting complex. Genes Dev. 2001;15(18):2396–2407.

44. Tian W, Li B, Warrington R, Tomchick DR, Yu H, Luo X. Structural analysis of human Cdc20 supports multisite degron recognition by APC/C. Proc Natl Acad Sci. 2012;109(45):18419–18424.

45. Thompson MJ, Jiggins CD. Supergenes and their role in evolution. Heredity. 2014;113(1):1–8.

46. Clarke CA, Sheppard PM. Super-genes and mimicry. Heredity. 1960;14:175–185.

47. Aulchenko YS, Ripke S, Isaacs A, van Duijn CM. GenABEL: an R library for genome-wide association analysis. Bioinforma Oxf Engl. 2007;23(10):1294–1296.

48. Supple MA, Hines HM, Dasmahapatra KK, Lewis JJ, Nielsen DM, Lavoie C, et al. Genomic architecture of adaptive color pattern divergence and convergence in Heliconius butterflies. Genome Res. 2013;23(8):1248–1257.

49. Martin SH, Dasmahapatra KK, Nadeau NJ, Salazar C, Walters JR, Simpson F, et al. Genome-wide evidence for speciation with gene flow in Heliconius butterflies. Genome Res. 2013;23(11):1817–1828.

50. Benjamini Y, Hochberg Y. Controlling the False Discovery Rate: A Practical and Powerful Approach to Multiple Testing. J R Stat Soc Ser B Methodol. 1995;57(1):289–300.

51. Lunter G, Goodson M. Stampy: A statistical algorithm for sensitive and fast mapping of Illumina sequence reads. Genome Res. 2011;21(6):936–939.

52. Li B, Dewey CN. RSEM: accurate transcript quantification from RNA-Seq data with or without a reference genome. BMC Bioinformatics. 2011;12(1):323.

53. Anders S, Huber W. Differential expression analysis for sequence count data. Genome Biol. 2010;11(10):1–12.

54. Hardcastle TJ, Kelly KA. baySeq: Empirical Bayesian methods for identifying differential expression in sequence count data. BMC Bioinformatics. 2010;11(1):422.

55. R Development Core Team. R: A language and environment for statistical computing [Internet]. Vienna, Austria: R Foundation for Statistical Computing; 2011. Available from: http://www.R-project.org/

56. Van’t Hof AE, Nguyen P, Dalíková M, Edmonds N, Marec F, Saccheri IJ. Linkage map of the peppered moth, Biston betularia (Lepidoptera, Geometridae): a model of industrial melanism. Heredity. 2013;110(3):283–95.

